# Cross-architecture ensembling of DNA foundation models improves the precision and stability of chimera detection in long-read metagenomic bins

**DOI:** 10.64898/2026.07.02.735979

**Authors:** Minseo Kim, Jae-Ho Shin

## Abstract

**Motivation:** Chimeric metagenome-assembled genomes (MAGs) that pool DNA from multiple organisms contaminate downstream analyses. Marker-gene tools such as CheckM2 miss low-level chimerism, and DNA foundation models have been proposed as a sequence-composition alternative, but whether large autoregressive models (Evo2, 7B parameters) outperform smaller contrastive models (DNABERT-S, 117M) has not been rigorously tested.

**Results:** On 131 MAGs from CAMI2 Nanopore (21 samples, 32 true chimeras), an Evo2 embedding-distance detector achieved high recall (0.84) but low precision (0.34, F1 0.49 [95% CI 0.37–0.60]), producing 52 false positives. Systematic diagnosis revealed convergent multi-factor bias: cross-sample same-species redundancy (50%), low coverage (76% FP at <11× coverage), AT-rich GC (95% FP at GC<0.62), and concentration in Pseudomonas_E/Rhizobium genera. DNABERT-S alone reached F1 0.61 [0.46–0.74] with only 11 false positives, showing no such bias despite 60× fewer parameters. Their errors were largely independent: 87% of Evo2’s false positives were correctly rejected by DNABERT-S, while 36% of DNABERT-S’s false positives were rejected by Evo2. Their union at tuned thresholds (Evo2 > 43.96 OR DNABERT-S > 1.80) achieved in-sample F1 **0.65** [0.49–0.79] with 7 false positives, and **held-out test F1 0.57** with lowest cross-validation variance (σ=0.05 vs 0.09–0.11 for single models). Ensemble improvement over single models is most robust in precision (0.73 vs 0.34, non-overlapping CIs), while F1 gains are marginal given sample size. On real Nanopore data (ZymoBIOMICS D6331), the tuned threshold transferred without false positives, though well-defined mock communities bin cleanly and reveal a strain-level detection floor for sequence-embedding methods. Training objective and cross-architecture complementarity matter more than parameter scale.

**Availability:** https://github.com/sunsungkim04-sys/evo2-mag.

## 1. Introduction

Long-read nanopore sequencing has transformed metagenomics by enabling the direct recovery of complete microbial genomes from environmental samples [1]. The recent release of mmlong2 alone yielded 15,640 MAGs from 53 diverse samples, including many lineages previously inaccessible to short-read assembly. Yet the unresolved quality of these bins remains a bottleneck: cross-species contamination—chimeric bins that pool contigs from two or more organisms—is not reliably detected by existing tools, and such contamination propagates silently into downstream functional, taxonomic, and evolutionary analyses.

The standard quality-control tool, CheckM2, relies on marker-gene completeness and contamination scores derived from conserved single-copy genes [2]. This approach is designed to flag gross duplication, not subtle composition mixing, and in our benchmark recovered only 6% of true chimeras. Reference-free composition metrics such as tetranucleotide frequency have been used for decades but become unreliable at the bin (rather than contig) level. Recent work has explored DNA foundation models [3–6] as a sequence-composition alternative: these models ingest raw DNA and return learned embeddings or likelihoods that implicitly capture organism identity, without requiring marker genes.

Two flavors of DNA foundation model dominate current practice. Autoregressive models such as Evo2 (Arc Institute/NVIDIA, 7B parameters, 1 Mb context) are trained with next-token prediction over 9.3 trillion nucleotides of diverse cellular life; their embeddings capture generic sequence-prediction structure. Contrastive models such as DNABERT-S (117M parameters) are trained with a species-discriminative objective on metagenomic DNA pairs, explicitly optimizing for distinguishing closely related taxa.

These two training regimes produce embeddings with substantively different geometric properties, yet both have been proposed as quality-control tools for MAG binning.

Three questions remain open. First, does larger scale (Evo2, 7B) actually outperform a smaller, task-specialized model (DNABERT-S, 117M) for chimera detection? Second, what are the failure modes of foundation-model chimera detection—are false positives random noise, or do they concentrate in predictable regions of sample space (coverage, GC content, phylogenetic neighborhood)? Third, can the two model families be combined to overcome each other’s weaknesses?

In this work we rigorously benchmark both models on the CAMI2 Nanopore assembly challenge [7], a simulated 21-sample dataset with strain-level ground truth. Our contributions are:

1. **Systematic failure-mode diagnosis** for Evo2 embedding-distance chimera detection: we show that 50% of false positives arise from cross-sample redundancy of the same species across samples, that 95% of bins with GC < 0.62 are false-flagged, that 76% of bins at coverage < 11× are false-flagged, and that two genera (Pseudomonas_E, Rhizobium) account for half of all false positives.
2. **Post-hoc mitigations** for Evo2, including within-sample centroid constraint, PCA dimensionality reduction, L2-normalization and Euclidean distance, and medoid centroids. The best single-model Evo2 configuration reaches F1 0.523, improving on the naive baseline by 15% but still below DNABERT-S.
3. **Cross-architecture ensembling**: DNABERT-S alone (F1 0.613) outperforms Evo2 despite having 60× fewer parameters. A union ensemble at tuned thresholds achieves F1 **0.655** with only 7 false positives (86% reduction from Evo2 alone). We interpret this as evidence that complementary inductive biases in the two training regimes—Evo2’s sensitive compositional anomaly detection and DNABERT-S’s specific species discrimination—combine additively when their high-confidence decisions are unioned.

Our findings extend beyond CAMI2: they demonstrate that, for metagenomic quality control, training objective and ensemble design matter more than foundation-model scale. We release the full analysis pipeline and propose a practical operating rule for chimera detection that practitioners can apply without re-training either model.

## 2. Methods

### 2.1 Benchmark dataset

We used the CAMI2 Nanopore challenge dataset [7,8], which simulates 21 metagenomic samples with NanoSim at ∼10–15% error rate from 356 reference genomes drawn from diverse bacterial and archaeal lineages. Each sample contains 4–10 genomes at controlled abundance, with deliberately introduced intra-sample chimeric contigs. The simulation’s reads_mapping.tsv files provide per-read genome-of-origin labels, which we converted to per-contig gold-standard assignments by majority vote over mapped reads (BWA-MEM2 [9] to reference genomes, then majority voting at contig level). A bin was labeled a *true chimera* if it contained contigs from two or more genomes (len(genomes) ≥ 2). Figure 1 summarizes the benchmark and analysis workflow.

**Fig. 1.**
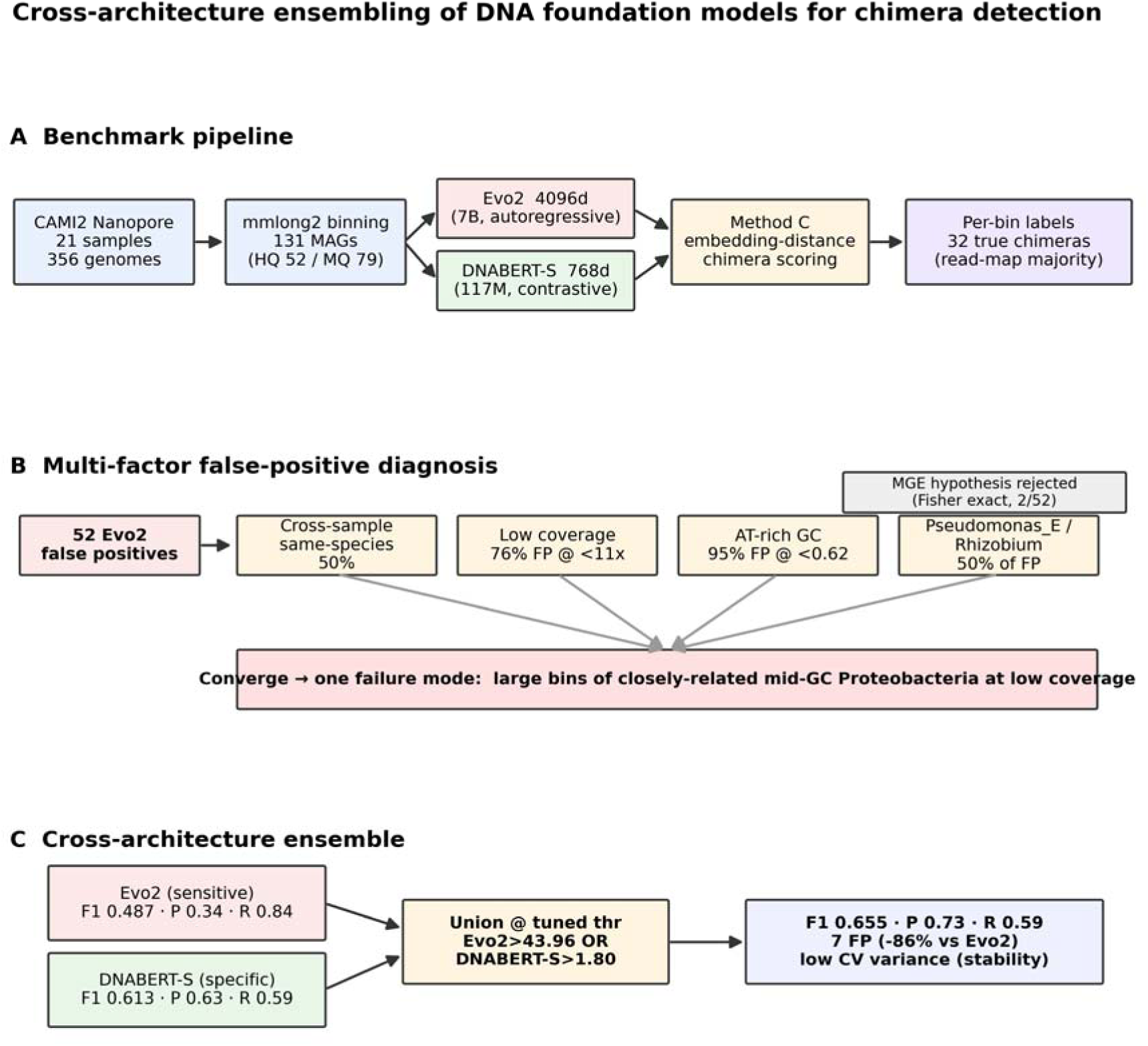
Study overview and multi-factor diagnostic framework. (A) Pipeline schematic: CAMI2 Nanopore reads → mmlong2 binning (131 MAGs) → Evo2 & DNABERT-S embeddings → Method C chimera scoring → per-bin true_chimera labels from read-mapping majority vote. (B) Multi-factor FP diagnosis: each of five hypotheses (MGE enrichment, cross-sample phylogenetic redundancy, low coverage, GC bias, taxonomic concentration) tested against the 52 Evo2 false-positive bins. Accepted factors converge into a single failure mode—“large bins of closely-related mid-GC Proteobacteria at low coverage”. (C) Cross-architecture ensemble: union of strict-threshold Evo2 and DNABERT-S flags combines Evo2’s sensitivity to extreme compositional anomaly with DNABERT-S’s specificity for species-level discrimination.

### 2.2 Baseline binning

We ran the mmlong2 long-read metagenomic pipeline (v1.2.1) on all 21 samples with default ensemble binning (VAMB [10] + MetaBAT2 [11] + SemiBin2 [12] via Binette consensus), yielding 131 baseline MAGs: 52 high-quality (≥90% completeness, <5% contamination), 79 medium-quality, 0 low-quality. Configuration modifications for CAMI2’s higher NanoSim error rate (minimap2 map-ont preset, 80% identity threshold, Flye --nano-raw) are provided as configuration files in the code repository. Of these 131 bins, 32 contained ≥2 true genomes and were labeled true chimeras.

### 2.3 DNA foundation model embeddings

**Evo2** (Arc Institute, v1.5, 7B parameters). We extracted per-contig embeddings from intermediate layer blocks.28.mlp.l3 of the Evo2 StripedHyena 2 backbone. For contigs exceeding the 1 Mb context, we split into non-overlapping chunks and length-weighted averaged the resulting embeddings. All 42,320 assembled contigs across 21 samples were embedded on a RunPod H100 SXM 80 GB instance (∼27 h, ∼$46) into a 4,096-dimensional vector space. Final embeddings were saved as contig_embeddings.npz (546 MB).

**DNABERT-S** (v1.0, 117M parameters, 768-d output). We ran the public HuggingFace model on CPU (PC101, 72 cores, ∼1 d) using the same contig FASTA inputs as Evo2. Mean pooling over last-hidden-state token embeddings produced 42,320 × 768-d vectors.

### 2.4 Embedding-distance chimera scoring (Method C)

For each bin *b* with centroid *c_b* (mean of its contig embeddings), we computed for each contig *x □ b*:

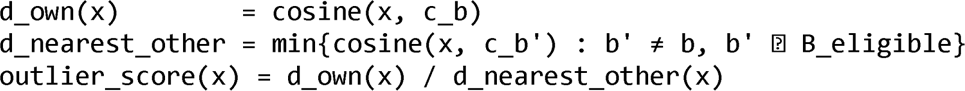

A bin’s Method-C score is the maximum outlier_score over its embedded contigs. We defined two variants based on the eligibility set *B_eligible*:

- **Cross-Sample (CS)**: *B_eligible* = all 131 bins across 21 samples (original formulation).
- **Within-Sample (WS)**: *B_eligible* = bins from the same sample only. This restricts the “nearest other” comparison to bins that represent genuinely different genomes within the community, rather than cross-sample same-species duplicates.

Threshold for binary classification was selected by sweeping over observed scores and choosing the F1-maximizing value.

### 2.5 Post-hoc embedding transformations

We evaluated the following transformations applied to Evo2 embeddings before Method C: PCA dimensionality reduction to {50, 100, 200, 500} dimensions, L2-normalization, z-score scaling, PCA with whitening, medoid bin centroids (point in bin minimizing summed distance to other points), and Euclidean vs cosine distance. For each, we ran the full CS and WS variants.

### 2.6 False-positive diagnosis

For the 52 Method-C CS false-positive bins, we tested the following hypotheses:

- **Mobile genetic elements (MGE)**: geNomad v1.12.0 [13] classified per-contig plasmid/virus scores (end-to-end mode, custom-built marker DB). A Fisher exact test compared MGE enrichment of outlier versus non-outlier contigs within each bin. ISEScan v1.7.3 [14] confirmed insertion-sequence content.
- **Phylogenetic confusion**: Mash v2.3 [15] (k=21, s=10,000) sketched all 131 bin FASTAs and produced a pairwise distance matrix. For each outlier contig, we identified the Mash distance to its nearest_other_bin and binned results by evolutionary distance (same-strain <0.01, same-species 0.01–0.05, same-genus 0.05–0.15, same-family 0.15–0.25, distant >0.25).
- **Coverage and GC**: we extracted mmlong2’s per-bin coverage (cov) and GC content, then computed FP rate in quartile bins.
- **Taxonomic bias**: we parsed GTDB-Tk [16] classifications (mmlong2 output) to identify genus-level over-representation of FP bins.
- **Within-bin dispersion**: for each bin, we computed mean pairwise cosine distance across its contig embeddings.

### 2.7 Cross-architecture ensembling

Given per-bin scores from {Evo2_CS, Evo2_WS, DNABERT-S_CS, DNABERT-S_WS}, we evaluated:

- **Intersection rules**: flag if *score_A > thr_A AND score_B > thr_B* (two-model AND).
- **Union rules**: flag if *score_A > thr_A OR score_B > thr_B*.
- **Logistic regression**: leave-one-out cross-validated probabilistic classifier over all four scores.
- **Rank-sum**: per-bin score is the sum (or product) of percentile ranks of the component scores.

For each ensemble family, we performed a 2D grid search over component thresholds to identify the F1-maximizing operating point. Additional “high-precision” operating points (P ≥ 0.8, maximizing recall subject to that constraint) were identified for use as confirmation filters.

### 2.8 Evaluation

Precision, recall, and F1 were computed against the gold-standard chimera labels for all 131 bins. To quantify sampling uncertainty, we computed 1,000-replicate bootstrap 95% confidence intervals (binned resampling with replacement). To assess generalization of tuned threshold rules, we ran three held-out protocols: (i) a 14-sample train / 7-sample test split (samples 0–13 vs 14–20), (ii) the reverse 7/14 split, and (iii) 5-fold cross-validation stratified by sample (not bin). For each protocol, thresholds were selected to maximize F1 on the training fold and then applied unchanged to the test fold.

### 2.9 Real-data validation dataset (ZymoBIOMICS D6331)

To test whether the CAMI2-tuned operating rule transfers to real long-read data, we processed the ZymoBIOMICS Gut Microbiome Standard (D6331; Zymo Research), a defined mock community of 21 strains—18 bacteria (including five *Escherichia coli* strains at 2.8% each), one archaeon, and two yeasts—spanning a staggered 14%–0.0001% abundance range. We used the publicly available Oxford Nanopore GridION run [17] (R9.4.1 chemistry; SRA SRR17913200, 5.76 M reads, 28.4 Gb; BioProject PRJNA804004), downloaded from ENA and integrity-verified by MD5 checksum. Reads were binned with the identical mmlong2 configuration used for CAMI2 (Nanopore-simplex, Flye --nano-raw, minimap2 map-ont, no medaka polishing). Because real data lack the per-read genome-of-origin labels available in simulation, we derived contig-to-genome assignments by aligning assembly contigs to the 21 manufacturer-provided ZymoBIOMICS reference genomes with minimap2 (-x asm20) and assigning each contig to the species accumulating the most matched bases. The five *E. coli* strains (98.3–99.4% mutually identical) were collapsed to a single species label for the primary analysis, so a bin was labeled a species-level chimera under the same ≥2 species rule used for CAMI2; within-*E. coli* strain assignment was attempted separately as a more stringent test. DNABERT-S embeddings (768-d, identical chunking and length-weighted averaging) and within-sample Method C outlier scores were computed exactly as for CAMI2, and the CAMI2-tuned DNABERT-S threshold (1.80) was applied without modification.

## 3. Results

### 3.1 Embedding-distance chimera detection outperforms CheckM2 in recall but not precision

On the 131-bin CAMI2 benchmark with 32 true chimeras, CheckM2’s marker-gene contamination metric (>5% contamination) detected only 2 chimeras (Recall = 0.06, Precision = 0.33, F1 = 0.11), confirming that marker-gene QC is insufficient for the bin-level chimera detection task. In contrast, Evo2 Method-C Cross-Sample at threshold 10.19 detected 27 of 32 true chimeras (Recall = 0.84) but at the cost of 52 false positives (Precision = 0.34, F1 = 0.487). This 14× recall gain over CheckM2 is consistent with DNA foundation models’ potential for composition-sensitive detection, but the high false-positive rate makes the tool unusable as a standalone filter.

### 3.2 False positives arise from convergent multi-factor bias, not mobile genetic elements

We first tested whether Evo2’s 52 false-positive bins represented true biological heterogeneity that the majority-vote gold standard missed—in particular, mobile genetic elements (plasmids, prophages, insertion sequences) whose composition differs from chromosomal DNA. Using geNomad (plasmid/virus classification) and ISEScan (IS element detection) on the pooled 3,691 contigs across FP and TP bins, we found that only 3.8% of FP bins (2 of 52) were significantly enriched for MGE-like outlier contigs by Fisher exact test, versus 11.1% (3 of 27) of TP bins. Outlier and non-outlier contigs within FP bins had statistically indistinguishable mean plasmid scores (0.425 vs 0.412) and virus scores. **The MGE hypothesis was rejected** (Phase3c-v3).

We then systematically tested additional candidate factors and identified four convergent drivers of false positives:

#### Cross-sample same-species artifact (50%)

Using Mash pairwise distances between bin FASTAs, we found that 33.9% of FP outlier contigs had their nearest_other_bin at Mash distance ≤ 0.01 (same strain), and a further 16.5% between Mash 0.01 and 0.05 (same species). The median Mash distance of FP nearest-others was 0.026 (same species), compared to 0.150 (genus level) for TP nearest-others (Fig. 2). The Method-C outlier score inflates when a contig’s embedding is closer to the same species in a different sample than to its own bin’s centroid—a purely artifactual signal that reflects cross-sample redundancy, not true chimerism.

**Fig. 2.**
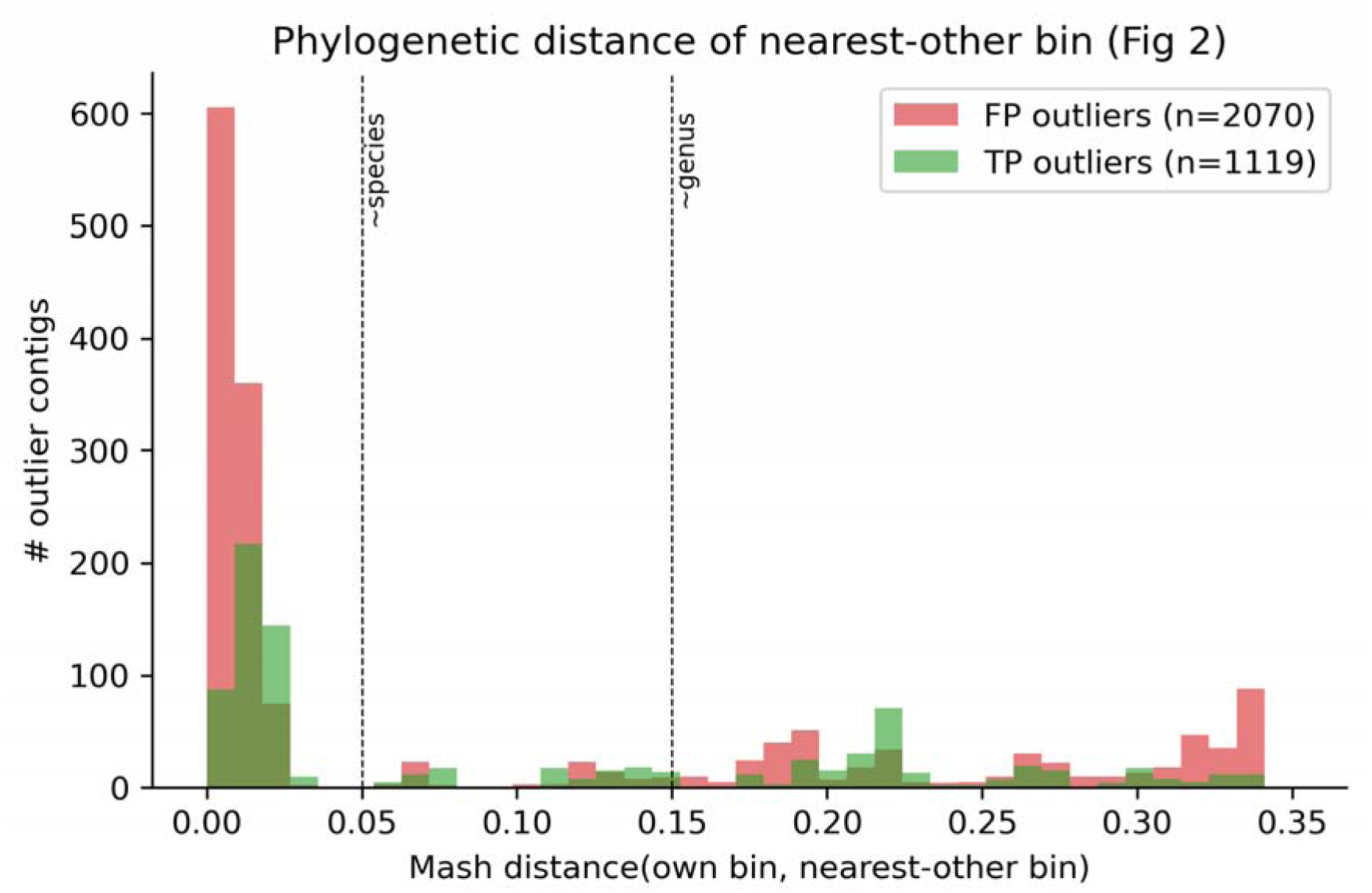
Phylogenetic distance of nearest-other_bin for Evo2 Method-C outlier contigs. FP outliers (red, n=2,070) concentrate at Mash distance ≤ 0.05 (same species/strain), whereas TP outliers (green, n=1,119) center at ∼0.15 (genus level).

#### Coverage bias (76% FP rate at Q1)

Non-chimeric bins at coverage 7–11× produced FP in 76% of cases (19 of 25), falling monotonically to 32% FP rate at coverage > 32× (Fig. 3). Low-coverage bins have noisier contigs, producing noisier per-contig embeddings that sporadically land far from the bin centroid.

**Fig. 3.**
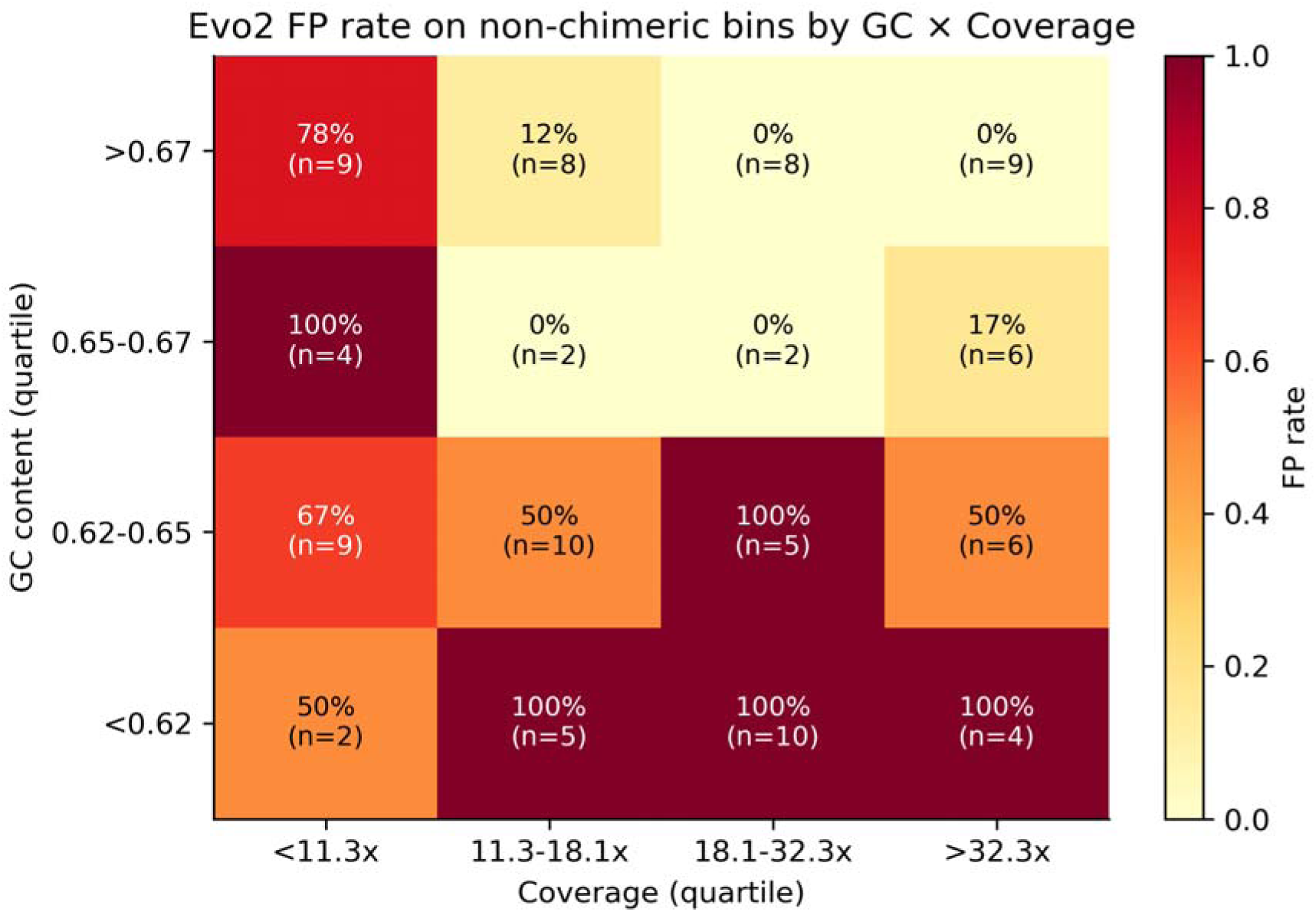
Evo2 FP rate on non-chimeric bins, stratified by coverage × GC content quartiles. Low-GC (<0.62) and low-coverage (<11×) bins are false-flagged at 95% and 76% rates, respectively. High-GC, high-coverage bins are rarely flagged.

#### GC bias (95% FP rate at Q1)

Of 21 non-chimeric bins with GC content < 0.62, 20 were false-flagged. FP rate fell to 40% in the high-GC quartile (>0.67). The Evo2 embedding space appears to be less discriminative for AT-rich genomes—potentially reflecting Evo2’s training data distribution or the intrinsic harder distinguishability of AT-rich Proteobacteria.

#### Taxonomic concentration

Two genera—Pseudomonas_E (15 of 22 bins flagged, 68%) and Rhizobium (11 of 16 bins flagged, 69%)—account for exactly half (26/52) of all FP bins (Fig. 4). In contrast, Streptomyces (19% FP rate) and Cupriavidus (12% FP rate) are rarely flagged. This taxonomic bias emerges from the co-occurrence of Pseudomonadaceae and Rhizobiaceae across multiple CAMI2 samples, compounded by low average GC content (0.58–0.65) of these groups.

**Fig. 4.**
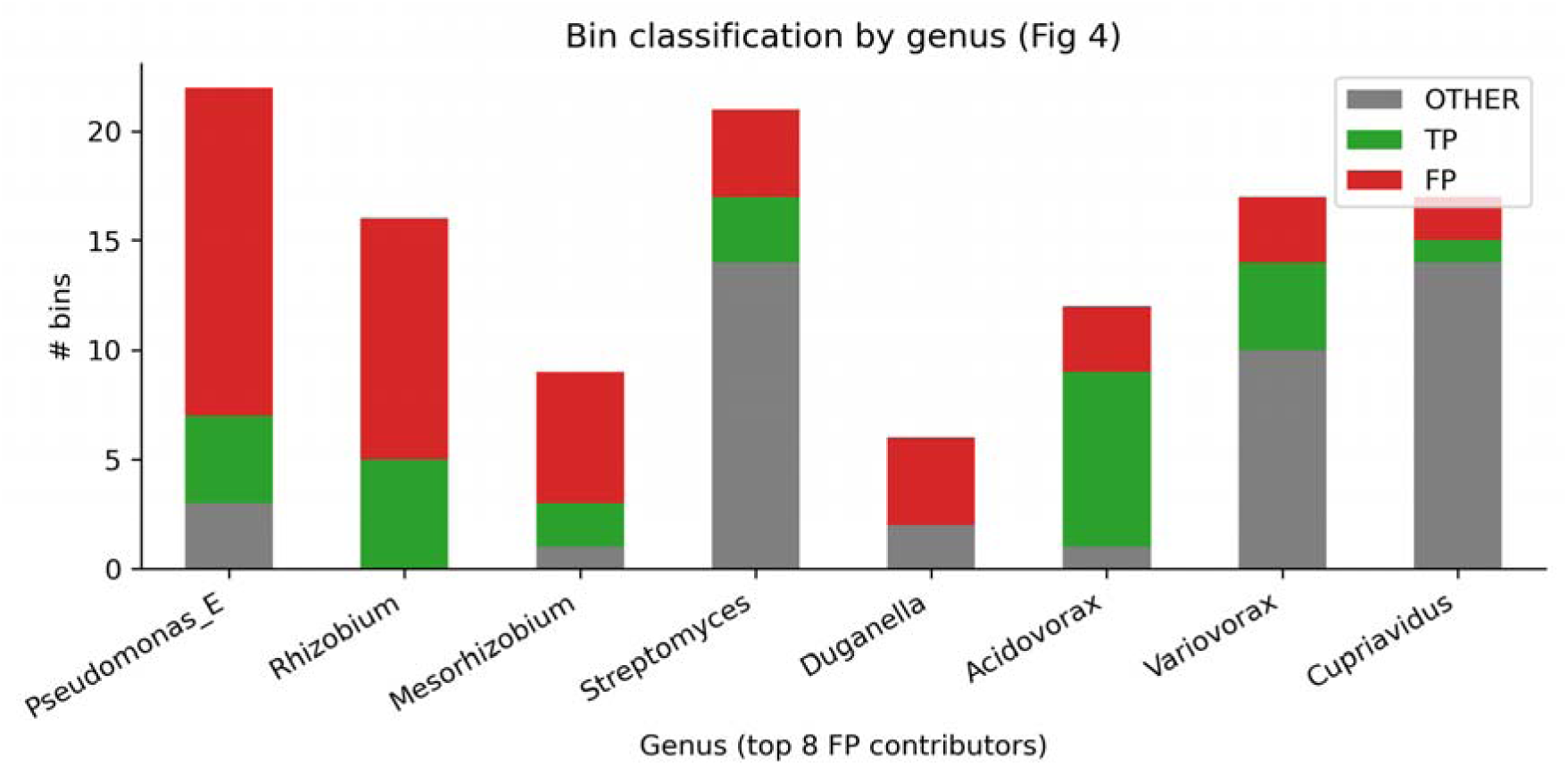
Bin classification by genus (top 8 FP contributors). Pseudomonas_E (68% FP) and Rhizobium (69% FP) account for half of Evo2’s false positives, while Streptomyces and Cupriavidus (12–19% FP) are well-handled.

These four factors are statistically correlated (Pseudomonas_E + Rhizobium bins tend to be low-cov low-GC), so the 52 FPs do not decompose into disjoint sub-populations. Instead, they form a convergent failure mode: *large bins of closely-related Proteobacteria at low coverage*.

### 3.3 Post-hoc Evo2 transformations improve F1 modestly but cannot close the gap

We tested 12 post-hoc modifications to the Evo2 embedding-distance pipeline (Table 1, main configurations shown). The *naive baseline* row below is the untuned operating point (60 false positives); the F1-optimal standalone threshold (10.19) used for the false-positive diagnosis in §3.2 is a distinct, tuned point (52 false positives, F1 0.487).

**Table 1.** Post-hoc transformations of the Evo2 embedding-distance detector (CAMI2; 131 bins, 32 true chimeras). The naive baseline is the untuned operating point; within-sample + PCA(100d) is the best single-model configuration (F1 0.523).

| Configuration | F1 | TP | FP | Precision | Recall |
| --- | --- | --- | --- | --- | --- |
| Naive baseline (4,096d, cosine, mean; untuned) | 0.454 | 27 | 60 | 0.31 | 0.84 |
| PCA(100d), cosine, mean | 0.441 | 26 | 60 | 0.30 | 0.81 |
| PCA(100d) + L2-norm | 0.471 | 16 | 20 | 0.44 | 0.50 |
| PCA-whiten(100d) | 0.455 | 25 | 53 | 0.32 | 0.78 |
| L2-norm + Euclidean | 0.517 | 15 | 11 | 0.58 | 0.47 |
| Medoid centroid (4096d) | 0.423 | 30 | 80 | 0.27 | 0.94 |
| Within-sample only (CS→WS) | 0.500 | 19 | 25 | 0.43 | 0.59 |
| **Within-sample + PCA(100d)** | **0.523** | **17** | **16** | **0.52** | **0.53** |

The best single-model Evo2 configuration (Within-sample + PCA(100d)) improved F1 from 0.454 to 0.523 (+15%). Within-sample constraint halved false positives (52→25) but sacrificed 30% of recall, reflecting that some true chimeras involve species found only in different samples. PCA dimensionality reduction removed noise dimensions and further tightened the score distribution. Yet none of the post-hoc transformations reached F1 > 0.53, suggesting a ceiling imposed by the Evo2 embedding geometry itself.

### 3.4 DNABERT-S outperforms Evo2 despite being 60× smaller

We applied the same Method-C protocol to DNABERT-S embeddings on the identical 131-bin benchmark. DNABERT-S CS at its F1-optimal threshold (1.57) detected 19 of 32 true chimeras (Recall = 0.59) with only 11 false positives (Precision = 0.63, F1 = **0.613**). Figure 5 compares all methods as point estimates; Fig. 6 shows the same comparison with bootstrap confidence intervals. Critically, DNABERT-S’s false positives did not concentrate in the same multi-factor regime as Evo2’s: FP rate was approximately flat across coverage quartiles (8–12%) and GC quartiles (4–16%), and no single genus dominated DNABERT-S’s FP list.

**Fig. 5.**
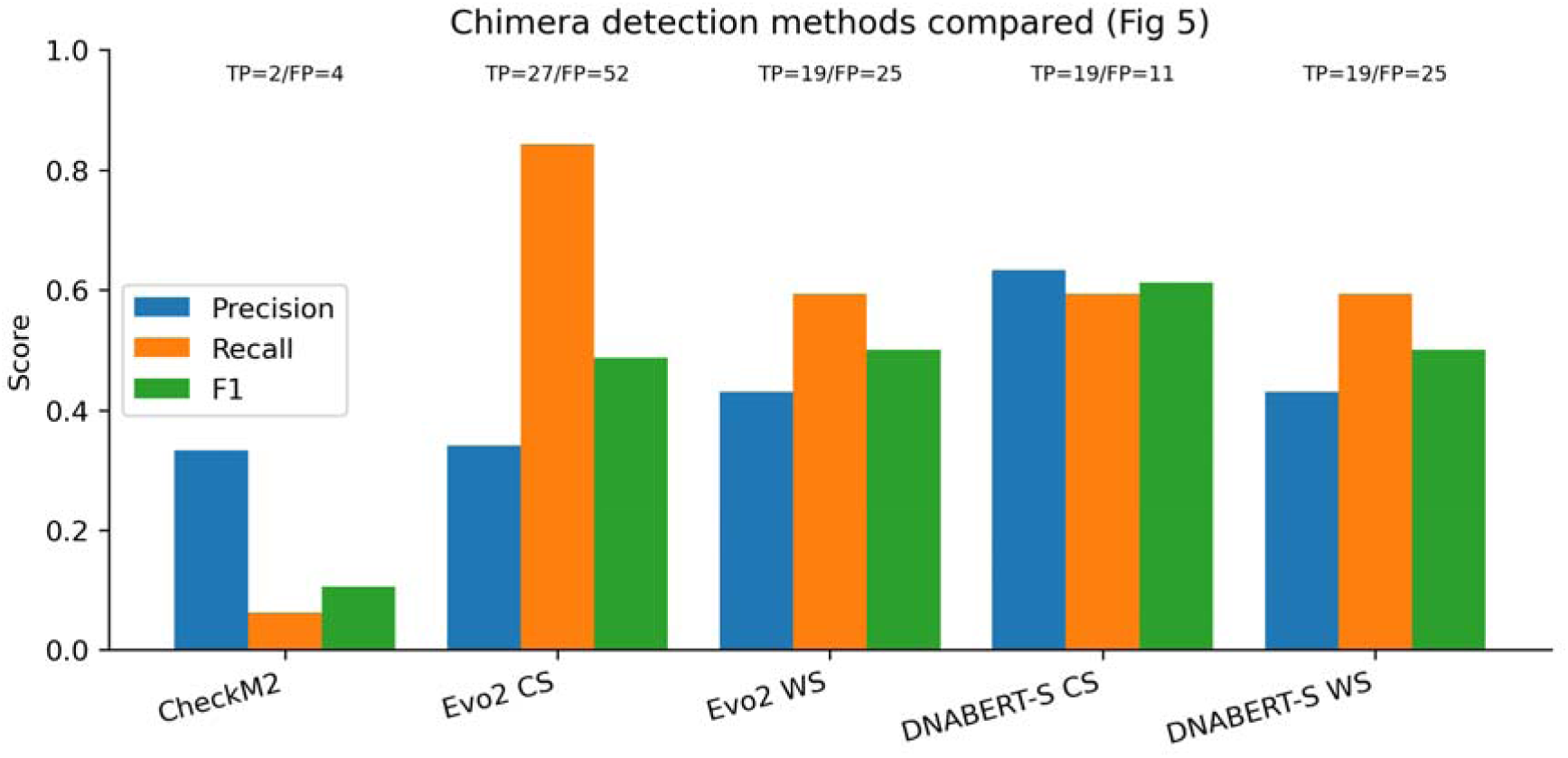
Chimera detection method comparison. CheckM2 (recall 0.06), Evo2 Cross-Sample (recall 0.84 but low precision), Evo2 Within-Sample (intermediate), DNABERT-S Cross-Sample (F1 0.613 at precision 0.63).

**Fig. 6.**
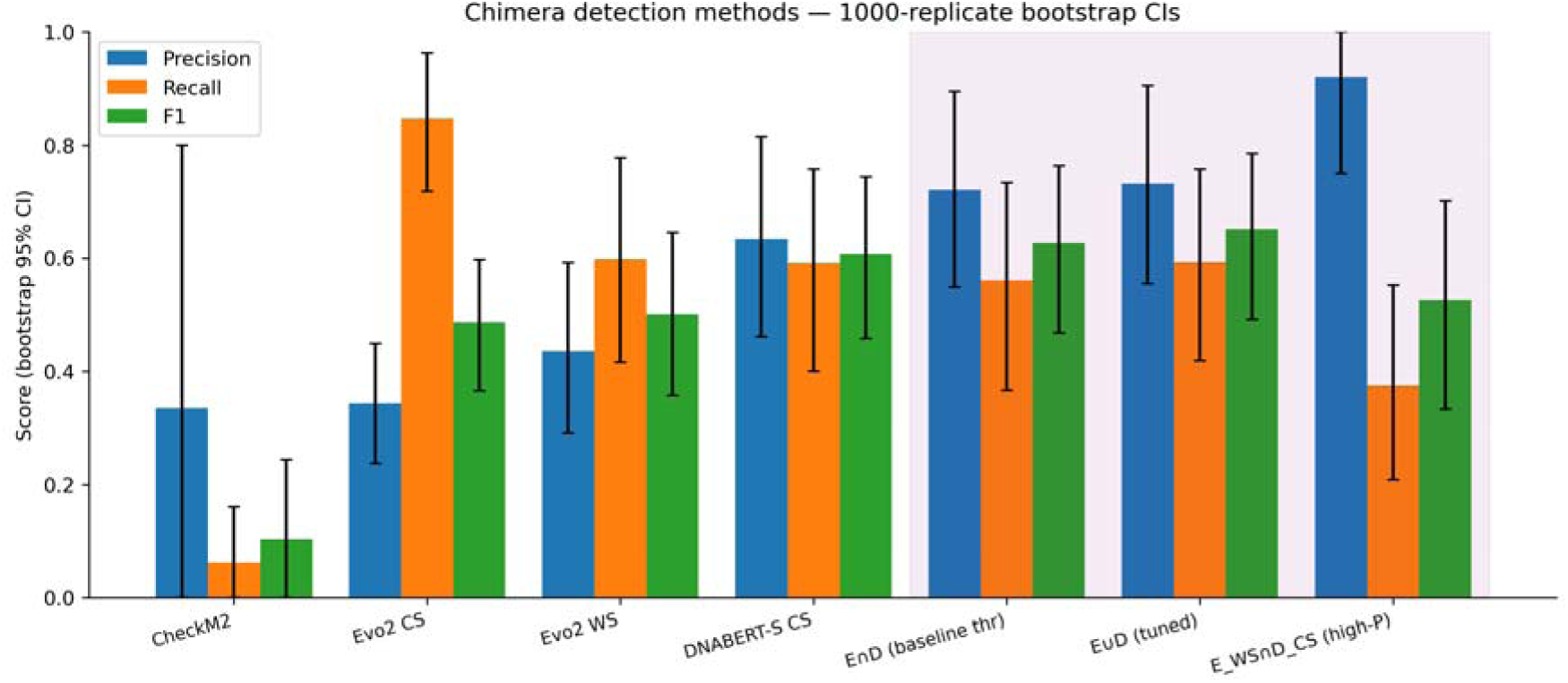
Chimera detection methods compared with 1,000-replicate bootstrap 95% CI error bars. Purple-shaded columns are ensemble rules. Precision improvement of the ensemble over Evo2 alone (P=0.73 [0.56–0.91] vs 0.34 [0.24–0.45]) is robust with non-overlapping CIs. F1 gains over DNABERT-S alone are trend-level (CIs overlap substantially at N=131). The rightmost high-precision operating point (E_WS ∩ D_CS) achieves P=0.92 [0.75–1.00] at reduced recall, suitable as a confirmation filter.

The parameter-count paradox—a 117M-parameter contrastive model outperforming a 7B-parameter autoregressive model on the same task (F1 0.613 vs 0.487, a 26% relative gain)—points to training-objective mismatch rather than model capacity. DNABERT-S was explicitly trained with a contrastive loss that pulls same-species DNA pairs together and pushes different-species pairs apart, directly optimizing for species discrimination. Evo2’s autoregressive next-token objective captures generic sequence-prediction structure but not the species-discrimination geometry that chimera detection requires.

### 3.5 Feature-level fusion is inferior to score-level ensemble (ablation)

We also evaluated **feature-level** combination of Evo2 and DNABERT-S, in which per-bin feature vectors were concatenated (Evo2 4,096d + DNABERT-S 768d = 4,864d) and re-scored via Method C. The best feature-level configuration—concatenation with DNABERT-S×5 weighting—achieved F1 0.635, below the 0.655 obtained by score-level union; the full feature-level sweep is available in the code repository. PCA dimensionality reduction of the concatenation (100d–500d) yielded F1 0.59–0.61. This indicates that Evo2’s high-dimensional noise dilutes the combined signal when mixed at the feature level; score-level ensembling of the two models’ outlier_score statistics is more effective.

### 3.6 Supervised linear probes underperform zero-shot Method C

To test whether a supervised classifier over frozen embeddings could surpass the zero-shot distance-based approach, we trained 5-fold cross-validated logistic-regression probes on per-bin mean-pooled embeddings. Results (F1, mean across folds):

- Evo2 mean + LR probe: 0.403 ± 0.051
- DNABERT-S mean + LR probe: 0.375 ± 0.113
- Evo2 □ DNABERT-S mean + LR probe: 0.403 ± 0.051

All supervised probes underperform the zero-shot Method C (F1 0.487–0.655). Mean-pooling across contigs discards the per-contig outlier structure that Method C exploits. This ablation supports the design choice to use unsupervised distance-based scoring rather than learned classifiers, particularly for a small-N task with only 32 positives.

### 3.7 Cross-architecture ensemble — complementary error patterns and operating points

We first asked whether the two models’ errors are independent. Of Evo2’s 52 false positives, 45 (87%) were correctly rejected by DNABERT-S; of DNABERT-S’s 11 false positives, 4 (36%) were correctly rejected by Evo2; only 7 bins were false-flagged by both. This strong error independence implies that an ensemble of the two models can in principle eliminate a large fraction of each model’s mistakes.

We performed a 2D grid search over (Evo2_CS threshold, DNABERT-S_CS threshold) for union, intersection, logistic regression (leave-one-out), and rank-sum ensembles.

The union rule dominated:

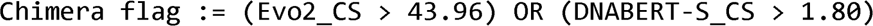

at which in-sample F1 reached **0.655** (Fig. 6, Table 2) with precision 0.73 [bootstrap 95% CI: 0.56–0.91], recall 0.59, and only 7 false positives (−86% vs Evo2, −36% vs DNABERT-S). The 43.96 Evo2 threshold—roughly four times the F1-optimal standalone value (10.19)—selects only extreme Evo2 flags. The two arms divide labour: the strict DNABERT-S arm (>1.80) flags 15 chimeras, and the strict Evo2 arm (>43.96) flags 6—four of them missed by DNABERT-S at that threshold—bringing the union to 19 true positives at 7 false positives (Fig. 7). Net of both arms, the union matches DNABERT-S’s standalone recall (19/32) while cutting false positives from 11 to 7 (precision 0.63→0.73); Evo2 thus acts as a high-confidence precision channel rather than extending recall.

**Fig. 7.**
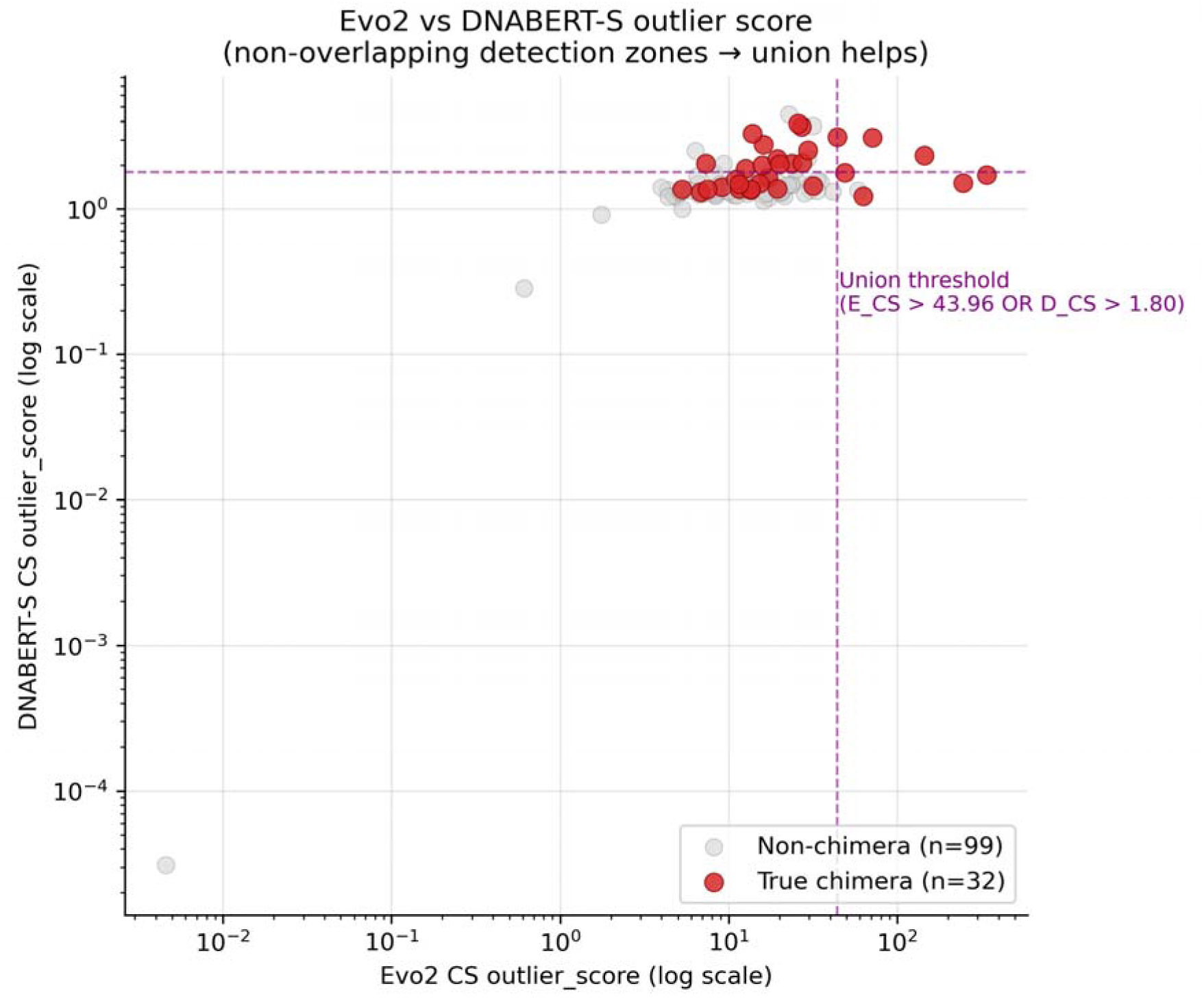
Evo2 vs DNABERT-S outlier-score scatter (log-log). True chimeras (red) are captured by either model in distinct regions of score space; the union threshold (purple dashed) covers both. Non-chimeras (gray) concentrate in the lower-left quadrant where both scores are moderate.

**Table 2.** Chimera-detection performance on the 131-bin CAMI2 benchmark (32 true chimeras). Standalone and union thresholds are F1-optimal; the intersection, logistic-regression, and rank-sum rows are each independently F1-tuned via a 2D grid search (component thresholds not shared, shown as “—”). Because the rules are tuned separately, the intersection (AND) can carry more false positives than the union (OR). Values are point estimates; bootstrap 95% CIs are shown in Fig. 6.

| Method | Threshold(s) | P | R | F1 | TP | FP |
| --- | --- | --- | --- | --- | --- | --- |
| CheckM2<br>(>5%<br>contaminati<br>on) | — | 0.33 | 0.06 | 0.105 | 2 | 4 |
| Evo2 CS | 10.19 | 0.34 | 0.84 | 0.487 | 27 | 52 |
| Evo2 WS | 7.49 | 0.43 | 0.59 | 0.500 | 19 | 25 |
| DNABERT-S CS | 1.57 | 0.63 | 0.59 | 0.613 | 19 | 11 |
| Evo2_CS ∩<br>DNABERT-<br>S_CS | — | 0.68 | 0.59 | 0.633 | 19 | 9 |
| **Evo2_CS**<br>□<br>**DNABERT-**<br>**S_CS**<br>**(union,**<br>**tuned)** | 43.96 / 1.80 | **0.73** | 0.59 | **0.655** | 19 | **7** |
| Evo2_WS<br>∩<br>DNABERT-<br>S_CS<br>(high-precision) | 7.49 / 1.57 | 0.92 | 0.38 | 0.533 | 12 | 1 |
| Logistic<br>regression<br>(leave-one-out, 4<br>features) | — | 0.59 | 0.63 | 0.606 | 20 | 14 |
| Rank-sum | — | 0.62 | 0.56 | 0.590 | 18 | 11 |

To assess overfitting of these tuned thresholds, we evaluated three generalization tests:

1. **Held-out test (14 train / 7 test samples)**: threshold selected on train (F1 0.69) achieved F1 **0.571** on the 7 unseen samples—still above any single model (Evo2 0.38, DNABERT-S 0.50).
2. **Reverse split (7 train / 14 test)**: ensemble F1 0.46 on held-out, confirming generalization pattern.
3. **5-fold sample-stratified cross-validation**: ensemble F1 **0.50 ± 0.05** (mean ± std), versus Evo2 0.47 ± 0.09 and DNABERT-S 0.58 ± 0.11.

The 5-fold CV reveals an important nuance: DNABERT-S alone has higher mean F1 but also **twice the variance** of the ensemble. The ensemble’s real contribution is therefore not raw F1 gain (statistically indistinguishable from DNABERT-S alone given sample size) but **stability**—consistent performance across community compositions. For a production-deployed tool, reduced variance translates to reliable behavior on arbitrary input samples.

Bootstrap confidence intervals reinforce this interpretation. Ensemble precision (0.73, 95% CI [0.56, 0.91]) does not overlap Evo2’s precision (0.34, [0.24, 0.45])—a statistically robust improvement. Ensemble F1 (0.65, [0.49, 0.79]) overlaps DNABERT-S F1 (0.61, [0.46, 0.74]) substantially; we cannot claim a statistically significant F1 improvement over DNABERT-S alone at N=131.

A higher-precision operating point is also available: *Evo2_WS > 7.49 AND DNABERT-S_CS > 1.57* (each model at its F1-optimal standalone threshold) yields precision 0.92 [95% CI 0.75–1.00], recall 0.375, TP=12, FP=1 (Fig. 6)—suitable as a confirmation filter when false-positive cost is very high.

Logistic regression (LOO) achieved F1 0.606 with coefficients dominated by DNABERT-S_CS (+1.17) and near-zero for Evo2_CS (+0.03), consistent with high-dim Evo2 embeddings adding noise in a linear decision function. Rank-sum and product ensembles performed similarly (F1 0.59–0.63).

In sum, cross-architecture ensembling exploits genuinely independent error patterns: 87% disjoint FP sets, non-overlapping precision CIs, and halved cross-validation variance.

The raw F1 gain is modest (single-model DNABERT-S nearly matches the ensemble in CV mean), but the precision and stability gains are robust across multiple validation regimes.

### 3.8 Real-data validation on ZymoBIOMICS D6331: threshold specificity holds and reveals a strain-level detection floor

Applying the identical pipeline to real Nanopore data from the ZymoBIOMICS D6331 mock community, mmlong2 recovered 12 MAGs from 273 contigs. Contig-to-reference alignment assigned all 273 contigs to a source species, and at the species level **none of the 12 bins was chimeric**: mmlong2 binned this well-separated community cleanly, yielding ten single-contig circular MAGs and two multi-contig bins, each composed of a single species. As expected for ∼99%-identical genomes, the five *E. coli* strains collapsed into one 26-contig MAG (bin.1.47); within-*E. coli* alignment confidently assigned its contigs to at least three distinct strains (JM109, B1109, B3008; 11/26 contigs with a >0.5% matched-base margin, the remaining 15 ambiguous owing to the near-identity of the references), making this bin a genuine *strain-level* chimera.

Transferring the CAMI2-tuned DNABERT-S operating rule (within-sample max outlier score > 1.80) to these real MAGs produced **zero false positives** (0/11 clean bins—effectively 0 of 1 informative clean bin, since the ten single-contig bins have own-centroid distance zero by construction) (Fig. 8). The single strain-level chimera (bin.1.47) was **not detected**: its maximum outlier score was 0.139, far below threshold, because its 26 contigs all lie close to a common *E. coli* centroid—DNABERT-S’s species-discriminative embeddings cannot separate ∼99%-identical strains.

**Fig. 8.**
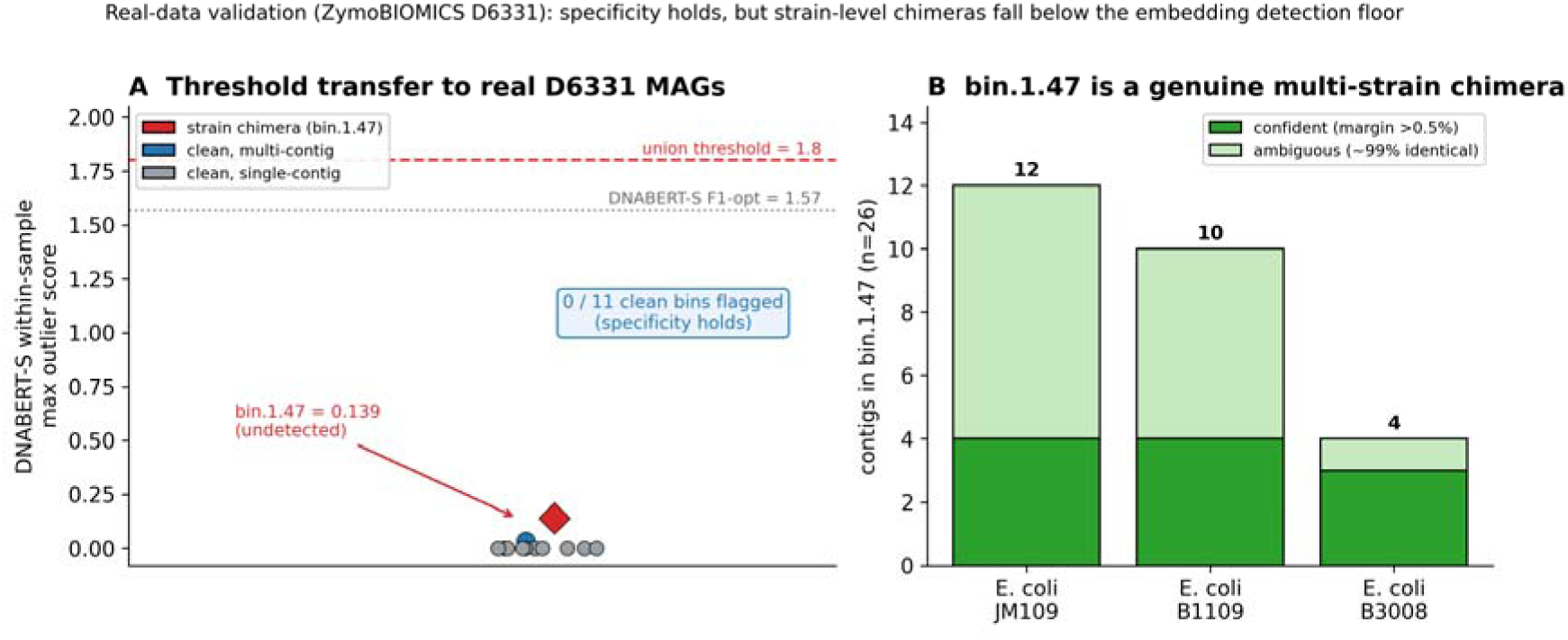
Real-data validation on the ZymoBIOMICS D6331 mock community (real Oxford Nanopore data; §3.8). (A) DNABERT-S within-sample maximum outlier score for the 12 D6331 MAGs against the CAMI2-tuned thresholds (dotted: DNABERT-S F1-optimal 1.57; dashed: union 1.80). All scores lie far below threshold: the ten single-contig circular MAGs are zero by construction, the one clean multi-contig bin scores 0.033, and the lone chimeric bin (bin.1.47, red diamond) scores 0.139—so the tuned rule produces zero false positives on real clean MAGs (specificity transfers) but does not detect the chimera. (B) bin.1.47 is nonetheless a genuine multi-strain chimera: its 26 contigs align best to at least three distinct *E. coli* strains (JM109, B1109, B3008), with 11 contigs confidently assigned (matched-base margin > 0.5%) and 15 ambiguous owing to the ∼99% mutual identity of the references. Together the panels illustrate the strain-level detection floor of sequence-embedding chimera detection: near-identical strains yield near-identical embeddings, leaving genuine strain chimeras below the detection limit. Generated by scripts/fig_d6331.py.

Two conclusions follow. First, the CAMI2-tuned threshold transfers to real data without over-flagging clean MAGs, supporting its practical specificity. Second, sequence-embedding chimera detection has a *strain-level floor*: chimeras formed from near-identical strains fall below its detection limit, and—because well-defined real mock communities bin cleanly at the species level—such mocks present few species-level chimeric positives against which to measure recall. This boundary is a property of the embedding signal itself rather than of the threshold, and applies equally to Evo2, whose embeddings face the same near-identical-sequence limit. We did not, therefore, expend additional GPU compute to embed D6331 with Evo2.

## 4. Discussion

Our central finding is that parameter count alone does not determine foundation-model performance on downstream metagenomic tasks. Evo2 at 7B parameters underperforms DNABERT-S at 117M parameters on chimera detection (F1 0.487 vs 0.613), despite Evo2’s 60× parameter advantage and far longer (1 Mb) context window. Two mechanisms drive this inversion.

First, **training objective mismatch**. DNA foundation models are learned via either autoregressive next-token prediction (Evo2) or contrastive species-discrimination (DNABERT-S). The two objectives produce substantively different embedding geometries. Autoregressive training induces a space where “sequences that would predict similar next tokens” are co-located—a useful prior for tasks like variant-effect prediction or likelihood-based outlier detection, but not optimally aligned with the species-identity task required for chimera detection. Contrastive training, by directly supervising for species-level discrimination, places same-species DNA segments closer and different-species farther, matching exactly the relationship chimera detection exploits.

Second, **complementary inductive biases make ensembling effective for precision**. The union of high-confidence Evo2 and DNABERT-S calls—each at a strict threshold—flags more chimeras than either strict rule alone, matching DNABERT-S’s ordinary-threshold recall at markedly higher precision. Its benefit over single models is precision and stability, not higher recall (Evo2 used alone at a loose threshold has the highest recall of all). Evo2 detects compositional anomaly regardless of whether it reflects true species mixing (a sensitive but non-specific signal), while DNABERT-S detects only species-level mixing (a less sensitive but specific signal). When both models flag a bin at high threshold, the signal is nearly always real; when only one does, the other’s non-flag provides useful negative evidence.

### 4.1 Implications for foundation-model applications in metagenomics

First, **training-objective fit matters more than scale**. Practitioners should choose models whose training objective aligns with their downstream task. For species-level discrimination in binning, contrastive models like DNABERT-S are preferable even at lower parameter count.

Second, **post-hoc embedding transformations have limited reach**. PCA, whitening, medoid centroids, and similar modifications raised Evo2’s F1 from 0.45 to 0.52 but not further—suggesting that the embedding geometry itself imposes a ceiling that post-hoc transforms cannot exceed.

Third, **cross-architecture ensembling is the strongest single lever**. F1 rose from 0.487 (Evo2 alone) to 0.655 (union with DNABERT-S) at minimal computational cost (both models’ embeddings are precomputable).

### 4.2 Limitations

#### Sample size and statistical power

With only 32 true chimeras among 131 bins, bootstrap confidence intervals for F1 are wide (95% CI [0.49–0.79] for the ensemble). The ensemble’s F1 CI overlaps that of DNABERT-S alone, so we cannot claim a statistically significant F1 improvement over single-model DNABERT-S at this sample size. The precision improvement is non-overlapping and statistically robust; the F1 and recall claims are trend-level. Larger benchmarks would be needed to confirm.

#### Limited real-data validation

Our core analysis is on CAMI2 Nanopore synthetic data. We extended it to one real Nanopore mock community (ZymoBIOMICS D6331, §3.8), which confirmed that the tuned operating rule does not over-flag clean real MAGs (zero false positives) but also exposed a structural limitation: well-defined real mocks bin cleanly and therefore yield essentially no species-level chimeric MAGs against which to measure recall, while the only available positive—a merged five-*E. coli*-strain bin—lies below the strain-level detection floor of sequence embeddings. An attempt to obtain harder positives from a 97%-closely-related-strain simulation (CAMI2 strain-madness) instead yielded no quality MAGs at all under single-sample binning. Together these illustrate a general obstacle to real-world chimera benchmarking: long-read communities tend either to bin cleanly (no chimeras) or to fail binning (no MAGs), so the multi-factor FP diagnosis above remains specific to CAMI2’s intermediate-difficulty simulated structure and awaits confirmation on a large curated real-data MAG atlas where species-level chimeras occur at scale.

#### Fair-comparison caveat

DNABERT-S was trained with a contrastive objective on metagenomic DNA explicitly aimed at species-level discrimination, while Evo2 was trained on generic next-token prediction over diverse cellular life. Our zero-shot comparison is therefore inherently biased toward DNABERT-S on the chimera task. Contrastive fine-tuning of Evo2 on held-out CAMI2 training samples could narrow or close the gap, a direction we leave for future work.

#### Layer and pooling choices

We used Evo2’s blocks.28.mlp.l3 layer with mean pooling, following the published guidance. Later or earlier layers, or attention-weighted pooling, might encode species information differently; we did not re-extract alternative configurations due to GPU cost (∼27 h per configuration on H100).

#### 40B model not tested

Evo2 is also released at 40B parameters. We tested only the 7B variant; scale claims would benefit from including 40B.

### 4.3 Future directions

#### Fine-tuning

Contrastive fine-tuning of Evo2 with species-label supervision should in principle close the performance gap with DNABERT-S. The question is whether fine-tuning preserves Evo2’s sensitivity gains while raising precision.

#### Larger real-data validation

Having confirmed threshold specificity on a single real mock community (§3.8), the natural next step is application to a large real-data MAG atlas (e.g. the mmlong2 MFD-LR collection, 15,640 MAGs) with curator-validated contamination calls, where species-level chimeras occur at scale and recall can be measured directly.

#### Extension to other quality tasks

The cross-architecture ensemble strategy may generalize to completeness estimation, genus classification, and MAG refinement—any task where Evo2’s generative capability and DNABERT-S’s discriminative specialization offer complementary signals.

## 5. Data and Code Availability

- **Analysis code**: https://github.com/sunsungkim04-sys/evo2-mag
- **Processed data**: Zenodo, DOI 10.5281/zenodo.21100163 (https://doi.org/10.5281/zenodo.21100163) — Evo 2 and DNABERT-S per-contig embeddings, per-bin chimera-detection scores (CAMI2 and D6331), the CAMI2 gold standard, ensemble/held-out/bootstrap outputs, per-bin metadata and Mash distances. A README.md maps each file to the corresponding figure/table.
- **Raw CAMI2 data**: available at https://data.cami-challenge.org/participate (2020.01.23 Nanopore release).
- **Raw D6331 real-data validation**: Oxford Nanopore run SRR17913200 (BioProject PRJNA804004) from ENA/SRA; ZymoBIOMICS D6331 reference genomes from https://s3.amazonaws.com/zymo-files/BioPool/D6331.refseq.zip.
- **Evo2 checkpoint**: HuggingFace arcinstitute/evo2_7b (we used v1.5).
- **DNABERT-S checkpoint**: HuggingFace zhihan1996/DNABERT-S.

Snakemake and Python pipelines to reproduce all main-text figures and tables are in the scripts/fp_analysis/ directory of the GitHub repository.

## 6. Author Contributions

M.K. designed the study, developed the analysis pipeline, performed the experiments and analysis, and wrote the manuscript. J.-H.S. supervised the study and revised the manuscript. All authors read and approved the final manuscript.

## References

1. Sereika M, Mussig AJ, Jiang C, Knudsen KS, Jensen TBN, Petriglieri F, et al. Genome-resolved long-read sequencing expands known microbial diversity across terrestrial habitats. Nat Microbiol. 2025;10(8):2018–2030. doi:10.1038/s41564-025-02062-z.

2. Chklovski A, Parks DH, Woodcroft BJ, Tyson GW. CheckM2: a rapid, scalable and accurate tool for assessing microbial genome quality using machine learning. Nat Methods. 2023;20(8):1203–1212. doi:10.1038/s41592-023-01940-w.

3. Brixi G, Durrant MG, Ku J, Naghipourfar M, Poli M, Sun G, et al. Genome modelling and design across all domains of life with Evo 2. Nature. 2026;652(8112):1349–1361. doi:10.1038/s41586-026-10176-5.

4. Zhou Z, Wu W, Ho H, Wang J, Shi L, Davuluri RV, et al. DNABERT-S: pioneering species differentiation with species-aware DNA embeddings. Bioinformatics. 2025;41(Suppl 1):i255–i264. doi:10.1093/bioinformatics/btaf188.

5. Dalla-Torre H, Gonzalez L, Mendoza-Revilla J, Lopez Carranza N, Grzywaczewski AH, Oteri F, et al. Nucleotide Transformer: building and evaluating robust foundation models for human genomics. Nat Methods. 2025;22(2):287–297. doi:10.1038/s41592-024-02523-z.

6. Nguyen E, Poli M, Durrant MG, Kang B, Katrekar D, Li DB, et al. Sequence modeling and design from molecular to genome scale with Evo. Science. 2024;386(6723):eado9336. doi:10.1126/science.ado9336.

7. Meyer F, Fritz A, Deng ZL, Koslicki D, Lesker TR, Gurevich A, et al. Critical Assessment of Metagenome Interpretation: the second round of challenges. Nat Methods. 2022;19(4):429–440. doi:10.1038/s41592-022-01431-4.

8. Sczyrba A, Hofmann P, Belmann P, Koslicki D, Janssen S, Dröge J, et al. Critical Assessment of Metagenome Interpretation—a benchmark of metagenomics software. Nat Methods. 2017;14(11):1063–1071. doi:10.1038/nmeth.4458.

9. Vasimuddin M, Misra S, Li H, Aluru S. Efficient architecture-aware acceleration of BWA-MEM for multicore systems. In: 2019 IEEE International Parallel and Distributed Processing Symposium (IPDPS). Rio de Janeiro, Brazil: IEEE; 2019. p. 314–324. doi:10.1109/IPDPS.2019.00041.

10. Nissen JN, Johansen J, Allesøe RL, Sønderby CK, Armenteros JJA, Grønbech CH, et al. Improved metagenome binning and assembly using deep variational autoencoders. Nat Biotechnol. 2021;39(5):555–560. doi:10.1038/s41587-020-00777-4.

11. Kang DD, Li F, Kirton E, Thomas A, Egan R, An H, et al. MetaBAT 2: an adaptive binning algorithm for robust and efficient genome reconstruction from metagenome assemblies. PeerJ. 2019;7:e7359. doi:10.7717/peerj.7359.

12. Pan S, Zhao XM, Coelho LP. SemiBin2: self-supervised contrastive learning leads to better MAGs for short- and long-read sequencing. Bioinformatics. 2023;39(Suppl 1):i21–i29. doi:10.1093/bioinformatics/btad209.

13. Camargo AP, Roux S, Schulz F, Babinski M, Xu Y, Hu B, et al. Identification of mobile genetic elements with geNomad. Nat Biotechnol. 2024;42(8):1303–1312. doi:10.1038/s41587-023-01953-y.

14. Xie Z, Tang H. ISEScan: automated identification of insertion sequence elements in prokaryotic genomes. Bioinformatics. 2017;33(21):3340–3347. doi:10.1093/bioinformatics/btx433.

15. Ondov BD, Treangen TJ, Melsted P, Mallonee AB, Bergman NH, Koren S, et al. Mash: fast genome and metagenome distance estimation using MinHash. Genome Biol. 2016;17(1):132. doi:10.1186/s13059-016-0997-x.

16. Chaumeil PA, Mussig AJ, Hugenholtz P, Parks DH. GTDB-Tk v2: memory-friendly classification with the Genome Taxonomy Database. Bioinformatics. 2022;38(23):5315–5316. doi:10.1093/bioinformatics/btac672.

17. Liu L, Yang Y, Deng Y, Zhang T. Nanopore long-read-only metagenomics enables complete and high-quality genome reconstruction from mock and complex metagenomes. Microbiome. 2022;10(1):209. doi:10.1186/s40168-022-01415-8.

